# A generalized quantitative antibody homeostasis model: regulation of B-cell development by BCR saturation and novel insights into bone marrow function

**DOI:** 10.1101/065730

**Authors:** József Prechl

## Abstract

In a pair of articles we present a generalized quantitative model for the homeostatic function of clonal humoral immune system. In this first paper we describe the cycles of B-cell expansion and differentiation driven by B-cell receptor engagement.

The fate of a B cell is determined by the signals it receives via its antigen receptor at any point of its lifetime. We express BCR engagement as a function of apparent affinity and free antigen concentration, using the range of 10^−14^ to 10^−3^ M for both factors. We assume that for keeping their BCR responsive B cells must maintain partial BCR saturation, which is a narrow region defined by [Ag]≈K_D_. To remain in this region, B cells respond to changes in [Ag] by proliferation or apoptosis and modulate K_D_ by changing BCR structure. We apply this framework to various niches of B-cell development, such as the bone marrow, blood, lymphoid follicles and germinal centers. We propose that clustered B cells in the bone marrow and in follicles present antigen to surrounding B cells by exposing antigen captured on complement and Fc receptors. The model suggests that antigen-dependent selection in the bone marrow results in 1) effector BI cells, which develop in blood as a consequence of the inexhaustible nature of soluble antigens, 2) memory cells that survive in antigen rich niches, identified as marginal zone B cells. Finally, the model implies that memory B cells could derive survival signals from abundant non-cognate antigens.

---

The enormous progress of bioinformatics, computation and mathematical modelling of biological phenomena is currently transforming all fields of biology, including immunology. The main intention of development of the model presented here is to provide a general quantitative framework for describing antibody-antigen interactions. General, because we attempt to insert all key developmental and differentiation events of B cells into the model in our first article, and all key soluble antibody-mediated antigen recognition phenomena in our second article. Quantitative, because we insert these events into a coordinate system clearly defined by concentration and affinity. While we try to address some questions about molecular mechanisms within the cells, effects of cytokines and chemokines, adhesion molecules, interactions with T cells, it is beyond the scope of the paper to answer those questions. Rather we focus strictly on interactions of surface or soluble immunoglobulins and antigens, yet being aware that B cells have several functions other than antibody production. The assumptions required for the development of the model place known phenomena in new perspectives and may also provide unexpected answers to existing questions.

## Application of the model to B-cell homeostasis

B cells are lymphocytes, “cells of the lymph”, which are present in the blood as part of the mononuclear cell fraction of white blood cells. They are produced in the bone marrow^1^ and are found throughout the body, reaching various tissues and organs via the blood and the lymphatics. B cells are defined by their ability to rearrange the genetic loci coding the surface immunoglobulin (sIg) of the B-cell antigen receptor (BCR) complex and by their ability to secrete antibodies in later stages of their development^2^. Surface or membrane Ig is composed of a heavy and a light chain that are linked by disulfide bonds in a L-H-H-L stoichiometry. The BCR complex contains in addition to the sIg various transmembrane and intracellular molecules that modulate signaling via the BCR^3^. This signaling is vital for all B cells from the moment of their commitment, since these signals drive survival, differentiation or death of the cell. Starting with the expression of the surrogate light chain B cells go through several cycles of activation, proliferation, survival, and antibody production, all governed by BCR engagement.

The generalized quantitative model (GQM) assumes that in order to deliver functional signals to the B cells the saturation of the BCR by antigen is regulated by adjusting the number of available cells and the apparent affinity of the interaction (Fig.1). Saturation is a function of the affinity of sIg and antigen and concentrations of these two. The concentration of potential antigens spans several orders of magnitude if we consider self-antigens with mM to pM concentration range in the blood^4^. External antigens may reach high concentrations at the site of entry and become diluted out by degradation or elimination. Complete absence of BCR engagement as well as the complete saturation of BCR would lead to inability to respond to changes (Fig.1). Theoretically the best sensitivity to changes in antigen concentration could be achieved by keeping saturation of BCR around 50%. This is the central tenet of GQM: B cells die, survive, proliferate and differentiate to keep their BCR partially saturated (Fig. 1 b and c). We shall use the concentration of free antigen [Ag] and apparent affinity, represented by the equilibrium dissociation constant K_D_, to map the fate of differentiating B cells. The advantage of using these two parameters instead of using BCR engagement alone is that key factors determining these events can be better separated. Concentration of free antigen refers to the antigen concentration after equilibrium has been reached with respect to binding to the sIg of the B cells in the examined compartment. Higher affinity interactions require less antigen for the same extent of BCR engagement, while low affinity interactions require more antigen (Fig.1). It is important to note that we shall be looking at compartments or niches: anatomical locations that are characterized by the presence of particular antigens, particular cells, particular molecules, all that in a defined and more or less confined space and volume. The ranges we use for the two dimensions reflect the following properties. B cells recognize and also discriminate antigen within an affinity range of K_D_=10^−6^–10^−10^ M^5^. We extended this range because beyond the kinetic properties of the interaction the composition of the immunological synapse, of the BCR itself, and the signalosome also influence the outcome of BCR engagement, resulting in different (higher or lower) apparent affinities. Thus, by apparent affinity we refer to the global strength of BCR engagement, determined by affinity and the aforementioned factors. The upper value of antigen concentration was chosen because the upper limit of concentration for macromolecules is in the millimolar range in molecularly crowded environments. In fact, the apparent affinity of interactions under conditions of molecular crowding, like an immunological synapse, may be higher than that observed dilute conditions^6^. The lower value shown represents 10 antigen molecules in 10 nanoliters, which we arbitrarily chose as a volume potentially sampled by a single B cell.

**Figure 1.**
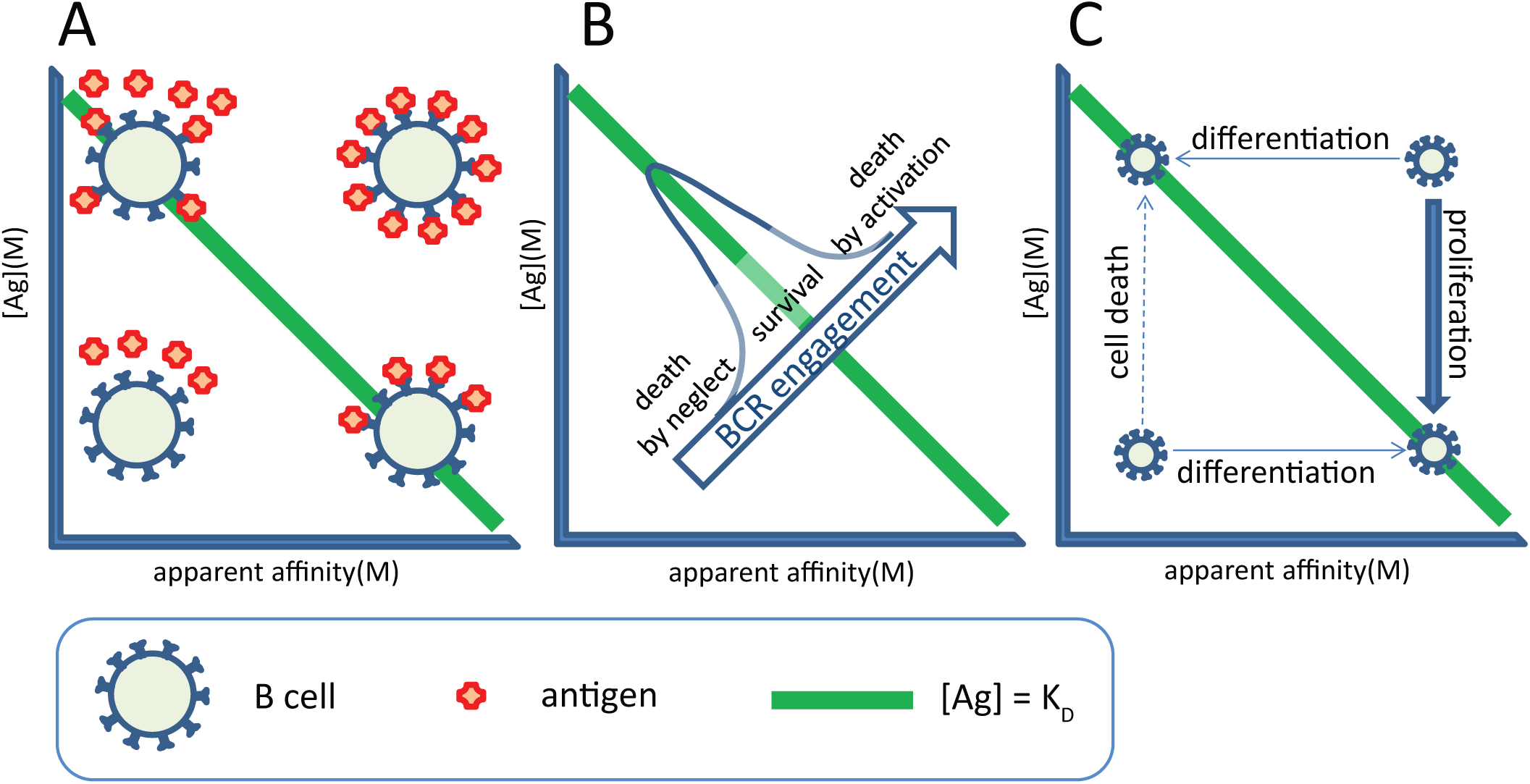
Outlines of the GQM for BCR-mediated B-cell regulation. A, Identical BCR engagement due to interactions with different apparent affinity is reached at different antigen concentrations. B, BCR engagement, as determined by apparent affinity and free antigen concentration at equilibrium, regulates B-cell survival. Green line indicates the comfort zone of the cells. C, Strategies for survival at BCR under- and overengagement.

By substituting BCR for Ab in the general equation defining equilibrium dissociation constant K_D_:

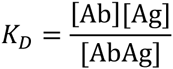

We derive

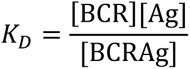

showing that K_D_=[Ag] when [BCR]=[BCRAg]. For the moment we neglect questions of avidity and shall regard [BCR] as the concentration of paratope and [Ag] as the concentration of epitope.

We assume that B cells have a “comfort zone” in our map, this comfort zone is defined by a region where [Ag] ≈ K_D_, enclosing the line representing 50% saturation of the BCR (Fig 1). Cells in the comfort zone adjust cell cycle to G0 phase, becoming resting, quiescent, long-lived cells. Such quiescent cells may receive tonic signals independent of antigen as well^7,8^. Activated, cycling B cells can return to this zone by controlling overall BCR numbers, simply achieved by their death or proliferation (vertical shifts in Fig.1). Cell death results in lower BCR numbers and released antigen increases [Ag], whereas proliferation leads to increased BCR numbers and a consequent decrease in [Ag] due to antigen binding to the cells and the limited local supply. Alternatively, cells can differentiate: change the real affinity of the interaction by generating new VH or VL domains and by mutating VH or VL domain sequences (horizontal shifts in Fig.1). Differentiation may involve apparent affinity changes when cells regulate BCR numbers and distribution on the cell surface, alter quality of signals delivered by the BCR itself or by its co-receptors (2^nd^ signals). These signals counteract cell death signals delivered via the Fas receptor family^9^. With these general considerations in mind we can start to take a look at the events of B-cell differentiation. We will be looking at mainly human physiology, in the different niches of B cell development, step-by-step.

## The bone marrow

B cells are generated from precursors in the bone marrow^1^. Hematopoetic stem cells give rise to pro-B cells, which are considered commited to becoming B cells once they express the transcription factor Pax-5^10^. Pre-BI cells rearrange their VDJ segments and enter our landscape by expressing a pre-BCR composed of a surrogate light chain (SLC) and a heavy chain. The SLC has been shown to bind to glycans^11^ or glycan binding lectins^12^ and thereby guide the recognition properties of the newly born pre-BCR. It is not known whether a ligand is required for the preBCR signaling but our GQM assumes so. Glycans abundantly present in the glycocalyx and membrane glycoproteins of cells are good candidate ligands. The glycocalyx displays self-identity beyond the genetic and proteomic information and can be considered the ultimate phenotype of the cell^13^ and is therefore a prime candidate for laying the foundations of the B-cell repertoire. Of note, analysis of serum IgM binding to different types of glycans revealed that blood group determines reactivity against glycans other than the blood group antigens^14^, pointing towards a genetically determined, glycan-mediated selection of precursor B cells. Selection requires the pairing of the rearranged heavy chain to SLC and potentially the ability to promote binding of the pre-BCR to its ligand. The SLC will thereby guide the pre-BCR by binding to molecules generally present on the cell surface. Successful binding translates to increased apparent affinity in our map (Fig. 2) and will trigger proliferation of the cells in order to return them to their comfort zone. Indeed cycling pre-BII cells, called large pre-BII, emerge at this point. These cells start rearranging their light chain VJ segments, leading to the expression of a BCR. Again we assume that the newly formed BCR has ligands in the microenvironment. Considering that proliferation is a sequence of repeated divisions, the resulting cells form clumps and are in contact with each other mostly. Could these newly formed cells themselves provide ligands for the pre-BCR or the BCR?

**Figure 2.**
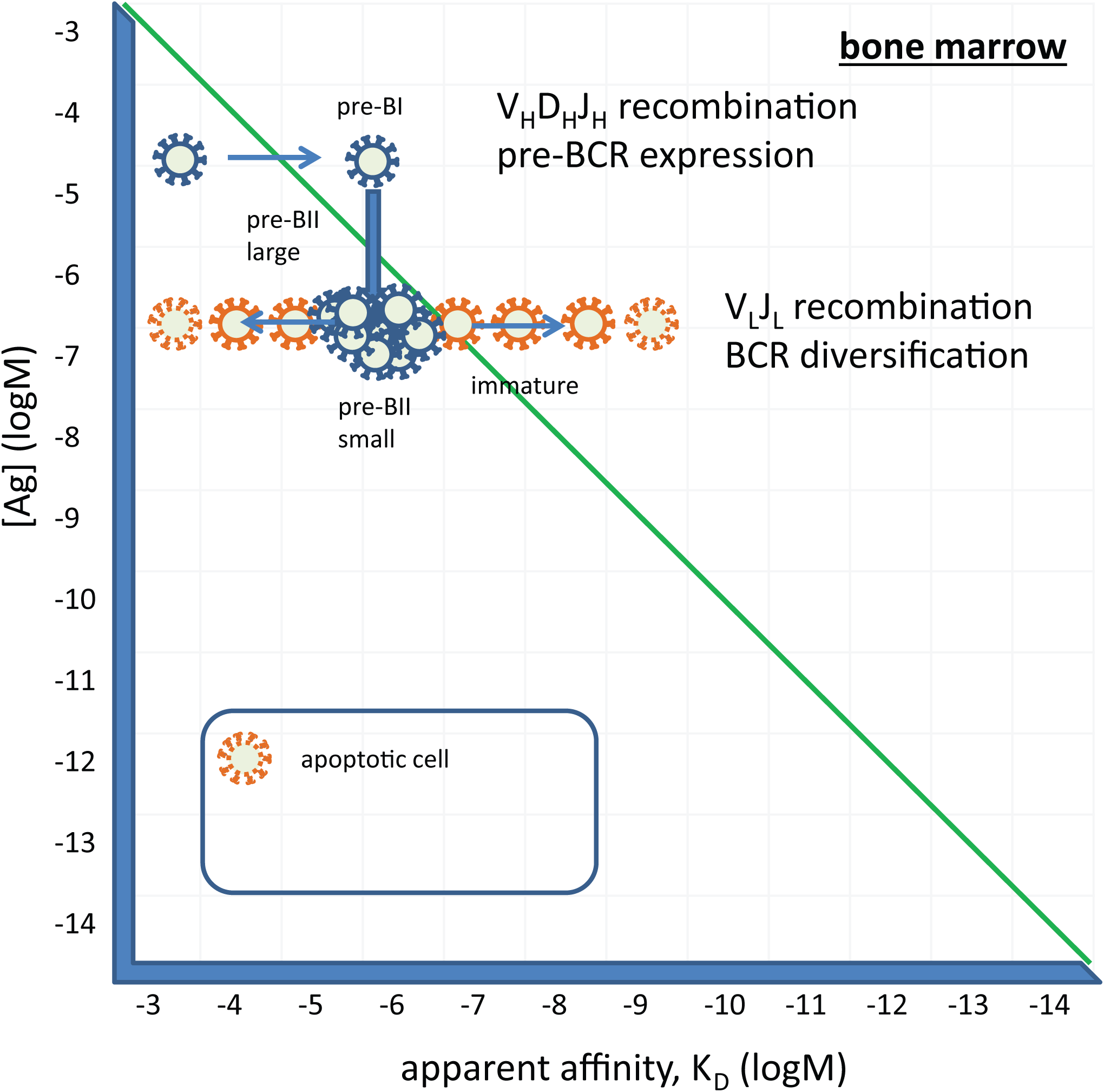
Development of B cells in the bone marrow. Pre-BI cells receive proliferation signals via the pre-BCR, entering several rounds of cells division. Cycling large pre-BII cells come to rest when they lose SLC, and start rearranging their light chains. Pre-BI small cells forming a signaling fit BCR reach their comfort zone and survive as immature B cells, other rearrange another light chain locus or die. Immature B cells leave the bone marrow.

One of the markers of early B cell forms in the bone marrow is CD93, a receptor for the collagen-like stalk of C1q. C1q binds to all sorts of molecules, recognizing charge and structure^15^. It also efficiently binds to IgM in immune complexes. The recently described IgM receptor, FcmuR, is also expressed by CD19 positive precursor B cells in human bone marrow^16^. Furthermore in the mouse, FcmuR has been shown to be required for B-cell development^17^. A simple – and by Occam's razor most likely - possibility to provide a feedback to newly forming IgM producing precursor cells is to display antigenic target molecules on the cells themselves. We suggest that **pre-BI and pre-BII cells present antigens by capturing antigen-bound C1q and IgM via their respective receptors**. These captured C1q and IgM molecules can thus provide their bound antigens as ligands for the BCR and mediate selection of the emerging repertoire. This proposition does not actually contradict the observed cell-autonomous pre-BCR signaling^18^, since no other cell type is required for signaling, pre-BI cells can stimulate each other via glycans displayed on the cell surface.

Selection events have long been associated with the modulation of self-reactivity. Developing B cells in the bone marrow are known to be deleted if they show aggressive self-binding. In other words **cells with low-affinity binding to self are selected here due to the fact that self-molecules are abundant** and these engage optimal number of BCR when the affinity of the interaction is low. With this in mind we can also accept that circulating IgM molecules – produced already during early stages of ontogenesis - with low affinity to self will do actually bind self-antigens because of the very high concentration of self-antigen. Thus, IgM molecules captured on the developing cells via FcmuR and/or C1qR are potential carriers self-molecules. These molecules can serve as ligands for the pre-BCR and BCR. Cells with inappropriate affinity to self will rearrange alternate chains (second VH or another kappa or lambda VH) until they reach their comfort zone or will die otherwise (Fig.2).

It is also important to contemplate on the diversity of the molecules displayed via C1q and IgM on the developing cells. Of the potential hundreds of different proteins, thousands of splice variants and glycoforms^4^ any and many may bind to these capture molecules. There is no antigen selection or antigen accumulation here, a great diversity of molecules can be displayed. Competing pre-BII small cells will therefore have to find epitopes that are shared by all these different antigens in order to receive survival signals via the BCR. The successful rearrangement of the light chain variable region, the successful paring of the light chain is now followed by a binding checkpoint^1^. **Selection pressures in the microenvironment of pre-BII cell favor polyspecificity** in the sense that binding to epitopes shared by the multitude of self-molecules or epitopes similar to each other on those molecules is an advantage. This flexibility is endowed by charge-dependent and hydrogen bonding interactions, which will allow association with the target but may not provide sufficient strength for stable binding, resulting in low-affinity interactions. Sequences of heavy chain hypervariable loops from germline antibodies are nearly optimal for such conformational diversity^19^.

Molecular abundance is thus defining factor for immunological self. Everything abundantly present during the development of the system is self immunologically. The developing immune system learns to identify all available molecules by generating a diversity of receptors and selecting those capable of recognizing this immunological self. Antigen-dependent selection in the bone marrow implies that while **in the embryonic and fetal stage of development immunological self is restricted to genetic self, after birth immunological self is extended to other molecules abundantly present in the organism**, which are not necessarily the products of self-genes. This results in a shift of characteristic autoreactivity of B cells observed before birth to a more modest self-reactivity after birth.

## The blood and extrafollicular sites of response

Immature B cells in the bone marrow enter the circulation. Upon arriving into the spleen these cells start changing their phenotype as described in the next section. But there are two cell types that cannot be properly accounted for in terms of differentiation, BI cells and marginal zone B cells. The former shows an activated phenotype, is dominant during embryonic development, is regarded as a source of natural antibodies, and responds to thymus independent antigens^20–22^. The latter is very well characterized by its anatomical location and is also responsive to TI antigens and to mitogenic stimuli in general^20,23^.

Blood is a unique antigenic environment being a fluid tissue. Antigens present in the blood plasma represent probably the whole proteome and glycome, with extreme variations in concentrations and extreme diversity of molecules^4^. Some of these molecules are dominantly present in blood, others are the result of leakage from tissues. Immature B cells entering the circulation will be suspended in this fluid tissue, testing their BCRs against nearly all possible molecules present in the body (Fig 3). They will react with these antigens with a range of affinities bringing a range of potential outcomes. A fraction of them will engage their BCR just to the extent of inducing modest cycling, in an attempt to reduce [Ag]. But unlike in solid tissues, where local antigen concentrations will decrease by increasing absolute BCR numbers, in a fluid with a vast volume of roughly 5 liters, antigen concentrations will not change at all. In a solid tissue cellular transport, diffusion rates, mechanical barriers slow down the replacement of antigen. In the blood the change in [Ag] due to a single cell division is negligible. Therefore these cells will keep replenishing themselves by remaining in an activated state (G1 phase of the cell cycle), dividing from time to time and adjusting their co-receptor set to be able to receive 2^nd^ signals for survival and differentiation. BI cells are thought to be a self-replenishing population with significant self-reactivity and responsiveness to danger signals^20^. We propose that **B1 cells develop from immature cells in the blood by being able to derive just the proper amount of survival and activation signals due to the combination of their apparent affinity and the amount of specific antigen present in blood** (Fig 3). Indeed B1 cells spontaneously produce antibodies which are presumably the source of natural antibodies^22^, as discussed in our accompanying paper (Prechl, submitted). Taking into account that the pre-BCR is selected for glycan reactivity it may not be surprising that antibodies secreted by B1 cells show preference for binding glycans^24^. In fact, autoreactivity of natural antibodies can be at least partially attributed to their glycan binding properties^24,25^.

**Figure 3.**
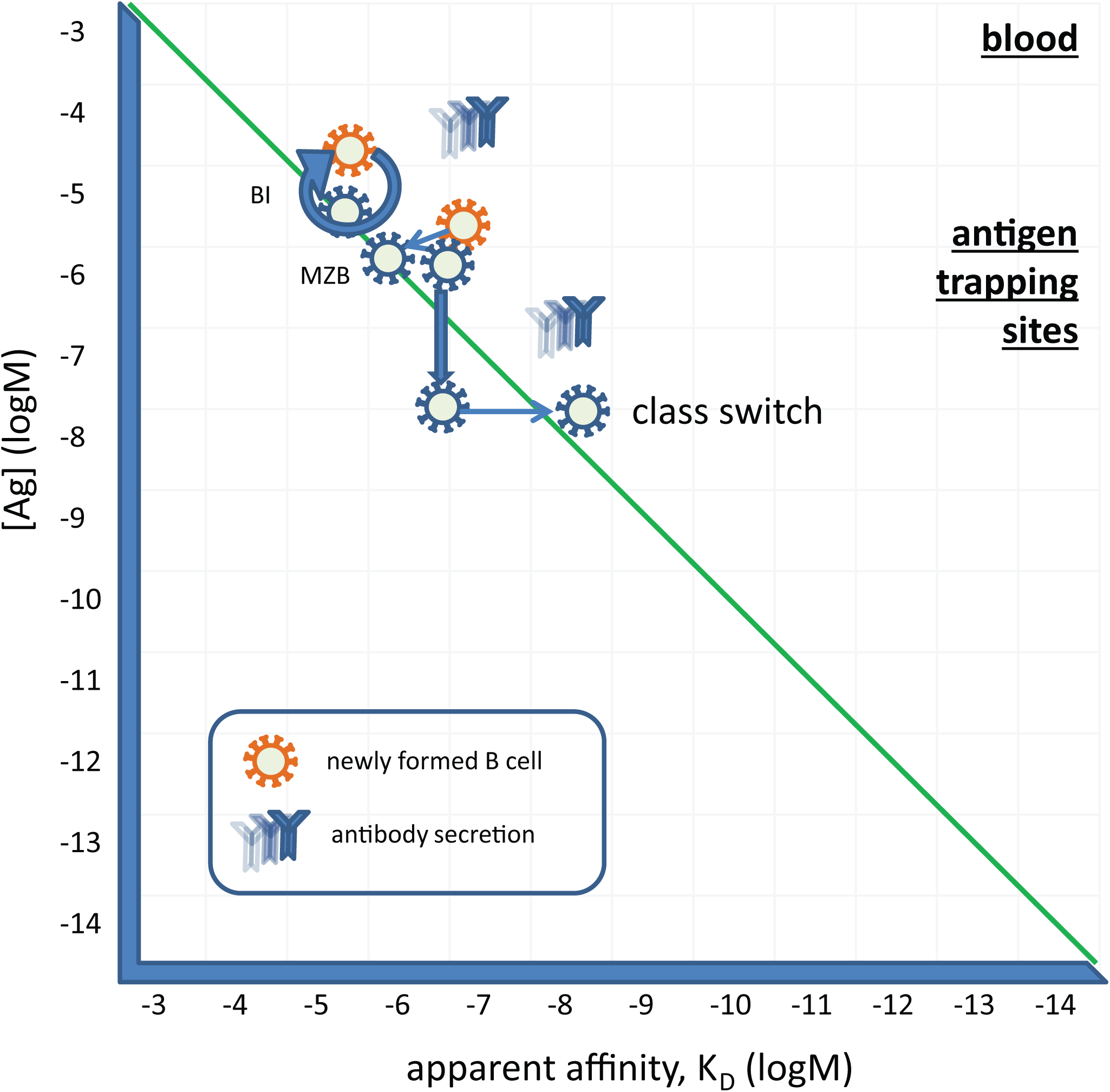
Differentiation of immature B cells in blood. Circulating B cells that receive moderate supraoptimal BCR signals may remain in a continuously activated state, dividing from time to time and secreting antibodies as BI cells. Otherwise these cells may differentiate into MZB cells and settle at antigen trapping sites, such as the marginal zone. Upon activation these cells may switch their heavy chain isotype in an attempt to increase apparent affinity of their BCR.

Another possible solution for cells showing strong reactivity to blood borne antigens is to modulate their BCR-derived signals. These cells may supplement BCR signaling with coreceptor and cytokine signals, and look for niches with adequate supply of their specific antigen. Development of marginal zone B cells relies on an important co-receptor, CD19^26^, and TLR signals may also contribute to their development^23^. These cells are found near sources of antigen: splenic marginal zone, subcapsular sinus in lymph nodes, tonsillar epithelium, subepithelial dome of Peyer's patches^23^. We therefore assume that these cells maintain their reactivity against immunological self and yet reach their comfort zone by modulating apparent BCR affinity. These features along with their responsiveness suggests that **marginal zone B cells represent a memory B cell population of the newly formed B cells.** Their specialized niches provide access of cells to a set of antigens necessary for their survival. These cells can be quickly and efficiently recruited for fast immune responses thanks to their location and reactivity.

A fraction of the cells egressing bone marrow may die due to neglect if their selecting antigens are present in the blood at lower concentrations. Other cells that encounter unknown antigens may engage their BCR with a higher apparent affinity, leading to activation and proliferation. Depending on the nature of the accompanying signals we can envisage several different outcomes. The simplest solution for the affected B cells is to divide as many times as required for reducing [Ag] and thereby reducing BCR engagement. This is accompanied by B cell blast formation and antibody production. This is the initiation phase of a primary immune response. In the presence of appropriate 2^nd^ signals (danger recognition receptors, btk signaling molecule, cytokines) cells may further proliferate and thus require adjustment of BCR engagement by changes to apparent affinity. A **solution for improving BCR apparent affinity is the switching of the isotype of the surface immunoglobulin molecule**, since class switched BCR show increased sensitivity to engagement^27^. This class switch is observed in the case of TI-3 responses, where myeloid cells provide contact and soluble signals for the activated B cell^28^. The resulting BCR will be of IgA, IgG or IgE isotype but with a low to medium affinity for the antigen (Fig.3).

Here we have to make a note on another function of the BCR, namely antigen uptake. High affinity and long lasting interactions with antigen trigger the internalization of the BCR complex leading to the transport of the antigen to the endolysosomal compartment^5^. Here the antigen is enzymatically degraded and peptides and lipids may bind to molecules of the major histocompatibility complex, MHC class II or CD1. Presentation of peptides and lipids to cognate helper CD4 T cells or NKT cells, respectively, provides a special signal for differentiation and gene expression. These responses are called TD-1 and TD-2, referring to the dependence of T cell help^28^. TD-2 responses lead to the formation of small germinal centers and minimal mutations. TD-1 responses are characterized by the development of germinal centers and extensive mutations that change not just the apparent but the true affinity of the BCR.

## Lymphoid follicles and the germinal center

Newly formed B cells, which based on their phenotype and function they are subdivided into transitional 1 and 2 cells (T1 and T2), enter the spleen. Transitional B cells acquire a different set of receptors capable of binding immune complexes: they lose C1qR and express CR2 instead^1^, becoming able to present complexes containing cleaved complement C3 products. Transitional B cells express BCR at high density rendering them sensitive to cell death but downregulate BCR expression upon becoming follicular BII cells. IgD expression is upregulated in follicular BII cells but disappears upon activation. Even though the exact role of surface IgD is not known, its characteristic and limited appearance in BII cells suggests it may have to do with providing survival signals for keeping these cells in the quiescent G0 state.

The diversity and concentration of antigens presented in the periphery, in addition to the presenting set of molecules, will largely depend on the environment of the host. Expression of FcmuR on the follicular B cells ensures that a set of antigens similar to those in the bone marrow will be displayed in the spleen at least, an organ that captures blood borne antigens just as the bone marrow. This is expected to provide a comfort zone of BCR signaling to the newly arriving T1 cells. The spleen is a likely place for the differentiation of transitional B cells into mature follicular BII cells. Surviving transitional B cells will become follicular BII cells at the end of the differentiation phase, having found their comfort zone of BCR engagement, and circulate in the body homing from follicle to follicle. Elsewhere, such as the lymph nodes and mucosa-associated follicles, self-antigens will be mostly supplemented by foreign, previously not encountered antigens.

We can compare the diversity of the molecules displayed in the follicle to that displayed in the bone marrow. Apart from the fact that the capture molecules themselves are different here, the principle is the same: feedback about the sampled anatomical region is provided by the molecules produced by activated cells of the repertoire. In the splenic follicles blood borne antigen is displayed, so the diversity of antigens may be similar to that in the bone marrow. In case a systemic antigen concentration becomes sufficiently high a more focused response may arise. Particulate antigens captured in the marginal zone may be transferred to the follicle, again favoring focused responses. Elsewhere in the periphery, such as the lymph nodes, Peyer's patches and other mucosal sites, particular antigens may become dominant, in the case of an injury or infection. Competing B cells will therefore have to increase their survival signals via the BCR in the presence of limited antigen. Thus, **selection pressures in the microenvironment of follicular B cells favor monospecificity**, cells acquiring optimal BCR signaling by changing the composition and/or the affinity of the BCR complex.

New non-self antigens entering a lymphoid follicle may react with BII cells with an unprecedented high affinity (Fig. 4). This interaction will trigger the proliferation of cells as discussed above, and also the kinesis of the cell, allowing it to move to the edge of the follicle and encounter T cells. Cognate interaction with a CD4 helper T cell will provide a 2^nd^ signal allowing survival and also the expression of genes that control somatic hypermutation(SHM). SHM is the process of generation of mutations mediated by enzymes involved in DNA repair processes^29^. The result of the process is the accumulation of mutations during repeated cell divisions and a diversified protein sequence coding for the BCR. Most mutations decrease the affinity of the interaction with antigen, these cells die during the selection process that follows. Because of the repeated divisions availability of antigen, as expressed by [Ag] will decrease, forcing the progeny of cells to compete. Cells with the highest affinity are therefore selected for cycling and longevity in the process (Fig. 4). High affinity binders may become effector cells and start secreting antibody, or otherwise modulate the affinity of their BCR and become memory cells. This process, called affinity maturation, increases the affinity of the BCR by replacing a flexible, conformationally diverse sequence with that of a more rigid conformation, showing better shape complementarity for the interaction with the targeted antigen^30,31^. An increased usage of amino acids conferring rigidity (proline) and filling space (histidine, phenylanaline) is observed at the antibody-antigen interface in matured human antibody sequences^32^. Increased affinity is endowed by a lock-and-key binding mechanism, which will allow slow dissociation from the target resulting in high-affinity interactions.

**Figure 4.**
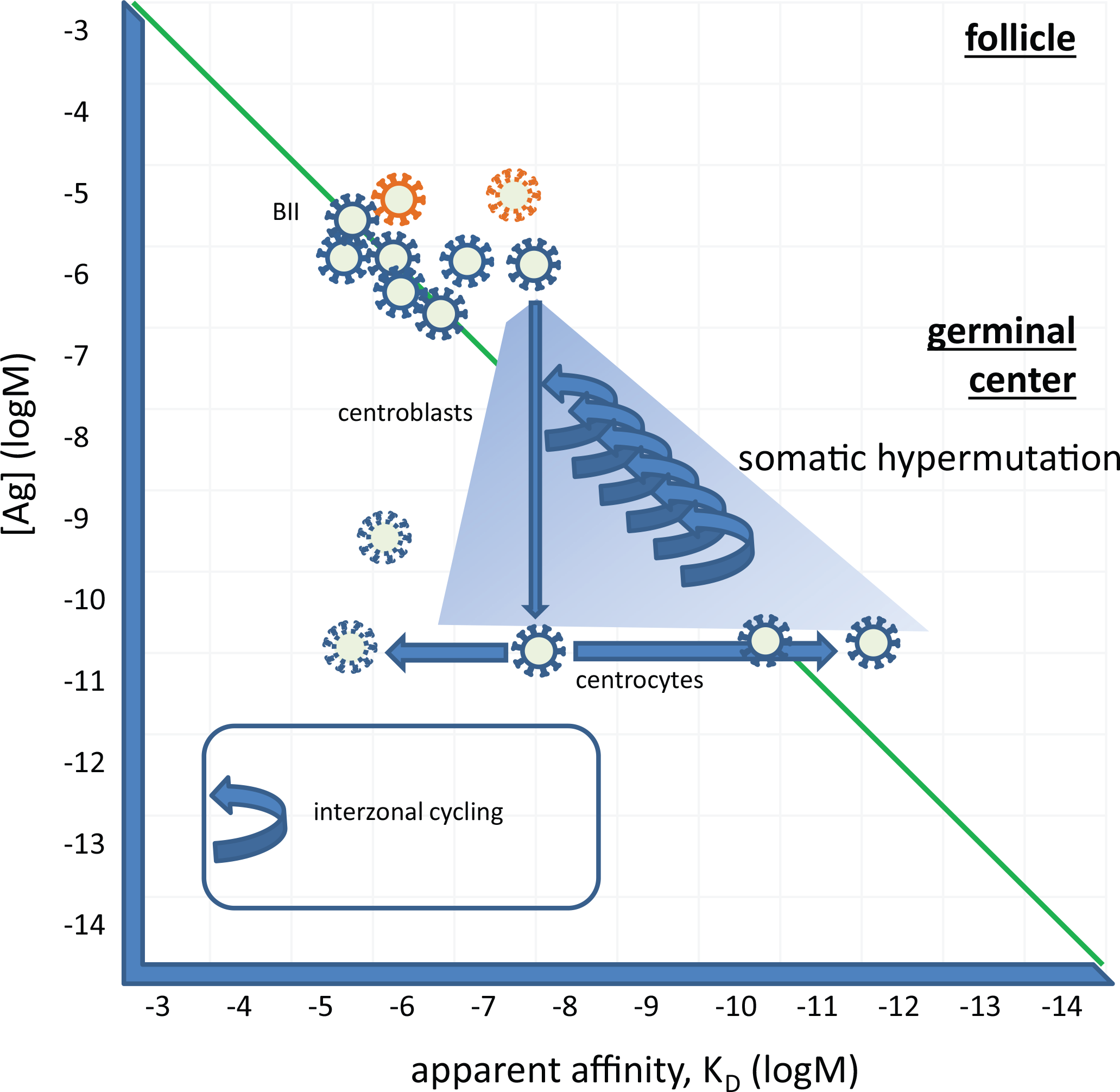
The generation of a germinal center. BII cells recirculate between follicles in a quiescent state. Here Ag from the blood, lymph, and mucosa is presented. When encountering an antigen that engages BCR they start dividing. Presentation of antigen via the MHCII to cognate Th cells provides costimulatory signals and results in the generation of a germinal center. Centroblasts devide and accumulate mutations, leading to changes in the affinity of the BCR. Competition favors BCRs with higher affinity. Selected high affinity cells return to the dark zone several times, capturing more antigen and starting another cycle of divisions.

The number of cell divisions in the germinal center has been shown to be regulated by the amount of antigen captured in the dark zone^33^, which corresponds to BCR engagement in our model. Higher affinity clones returning for antigen will capture more antigen in the same microenvironment with a given antigen concentration, triggering more rounds of division (Fig. 4). Heavy and light chain variable regions in affinity matured antibodies may contain up to 30 and 20 mutations, respectively^34^. Repeated vaccinations in humans led to an average affinity of 1.0 × 10^−9^ M, ranging from 1.0 × 10^−7^ M to 2.0 × 10^−12^ M^34^.

## Back to the bone marrow - memory generation and antibody secretion

According to our GQM B cells with strongly engaged BCR have two options to reach their comfort zone: modulate BCR signals or decrease [Ag] (Fig. 5). A potential solution for decreasing BCR engagement is to look for lower affinity interactions. With the advance of the immune response the eliciting antigen may become so scarce that it cannot provide survival signals any more. So some **cells may find niches with appropriate [Ag] but this antigen may not necessarily be the eliciting antigen itself**. These memory B cells^35^ can use antigen that is more abundant than the antigen that triggered selection of the cell. This interaction will be of lower affinity but still might keep the cell in the comfort zone in a resting quiescent state. The sites where antigen from different sources is most abundant should be close to the sources of self-antigen in blood, which is the extravascular sides of sinus endothelium in the spleen and bone marrow, the subcapsular sinus in lymph nodes, close to the tonsillar epithelium, and subepithelial dome of Peyer's patches – where MZ B cells are found as well. Analysis of the IgG+ memory cell repertoire found low-affinity self-reactivity of these cells^36^, in accordance with out model.

**Figure 5.**
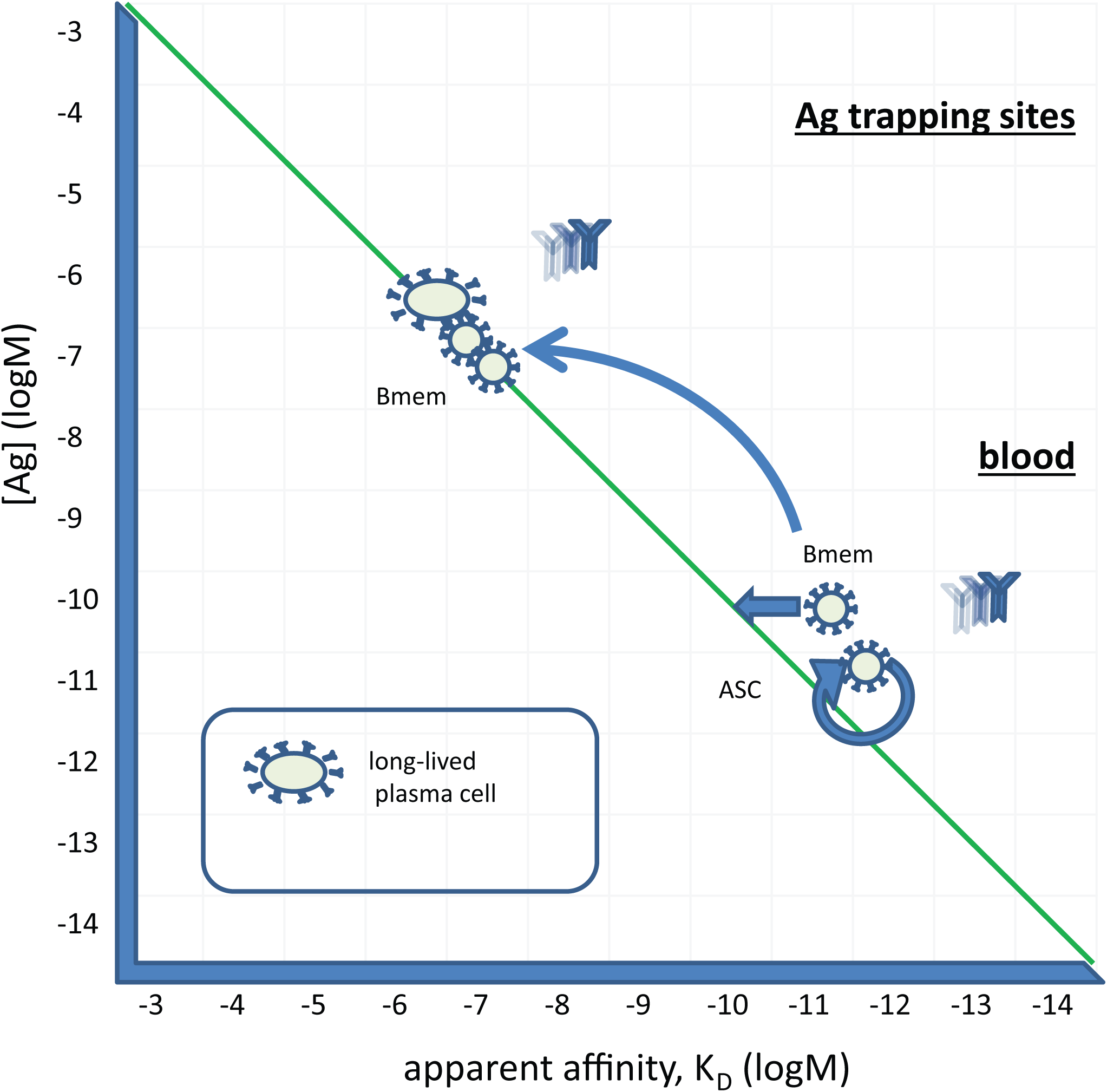
Generation of memory and antibody secreting cells. Cycling cells can reach their comfort zone by modulating their BCR signals and searching for a niche where a BCR ligand can provide survival signal. Until antigen concentration decreases these cells will continue to secrete antibodies and divide. Ideal sites for these cells, bone marrow perivascular niche, splenic marginal zone, lymph node subcapsular sinus, tonsillar epithelium, subepithelial dome of Peyer's patches are close to sources of antigen and are equipped with antigen presenting cells. Cells with similar BCR engagement may therefore reach two different states: memory by BCR signaling adjustment and ASC by cycling and exerting effector function.

So far we have only addressed the possibility reducing local [Ag] by doubling BCR numbers via cell division. B cells actually take steps to decrease global [Ag] by secreting a soluble form of the BCR, secreted antibodies. Secreted antibodies circulate in the blood and also reach extravascular sites where transport and permeability allows. Thus, circulating antibodies will cut the supply of antigen to the locale where B-cell response was initially triggered. Antibody secreting cells are also called plasmablasts and plasma cells, representing different stages of differentiation and rates and lengths of antibody production. How circulating antibodies fit into the GQM is described in our accompanying article.

## Summary and lookout

We approached the sequence of events leading to humoral immune response with the assumption that availability of antigen and affinity of the surface antibody-antigen interaction together define BCR engagement and B cell fate. In order to remain responsive to changes of antigen concentration in the microenvironment B cells proliferate, die and modulate BCR signaling properties. This assumption led to the proposal that B-cell selection in the bone marrow is antigen dependent. Antigen is transported to the bone marrow by immune complexes and displayed there on the developing B cells themselves. Obviously such a process needs to be initiated by antibody production, otherwise there would be no immune complex formation. So B cells are raised to identify self, perhaps first identified by C1q and other innate molecules, later expanding the range of recognized molecules in a recursive process where novel targets are brought for recognition with a continuously growing and changing antibody repertoire. Similar events take place in the secondary lymphoid organ follicles, where IgM-mediated antigen display is extended to antibodies with switched isotypes. The combination of cell expansion and differentiation events in the bone marrow, in the blood, in the follicles and germinal centers can be visualized in our map in a quantitative manner and confirm previous observations on antigen-driven responses.

The overall picture that emerges from the GQM is that there is an antigen-dependent, bone marrow-mediated and bone marrow-regulated homeostatic immune response to the immunological self, mediated mainly by B1 cells and MZ B cells. This is a dynamic, fluctuating immunity empowered by the supply of immature B cells selected for low affinity self-reactivity in the bone marrow. Selection events in the bone marrow in turn are modulated by antigens transported to the bone marrow via blood by antibodies secreted into the blood. Immune responses mediated by secreted antibodies are discussed in our accompanying paper on circulating antibodies and their network interactions.

## Acknowledgments

This paper is dedicated to the late Basel Institute for Immunology where I first learned about lymphocyte development and to Fritz Melchers who advised us to try to quantitate our microarray-generated antibody profiling data. I thank Dorottya Kövesdi for insightful discussions and suggestions and Gábor Koncz for critical assessment of the manuscript.

